# ABCal: a Python package for Author Bias Computation and Scientometric Plotting for Reviews and Meta-Analyses

**DOI:** 10.1101/2023.11.07.565991

**Authors:** L.S. Le Clercq

**Affiliations:** South African National Biodiversity Institute, Pretoria, 0001, South Africa; Department of Genetics, University of the Free State, Bloemfontein, 9300, South Africa

**Keywords:** Reviews, Authorship, Bias detection, Author bias, Scientometrics, Plots

## Abstract

Systematic reviews are critical summaries of the exiting literature on a given subject and, when combined with meta-analysis, provides a quantitative synthesis of evidence to direct and inform future research. Such reviews must, however, account for complex sources of between study heterogeneity and possible sources of bias, such as publication bias. This paper presents the methods and results of a research study using a newly developed software tool called ABCal (version 1.0.2) to compute and assess author bias in the literature, providing a quantitative measure for the possible effect of overrepresented authors introducing bias to the overall interpretation of the literature. ABCal includes a new metric referred to as author bias, which is a measure of potential biases per paper when the frequency or proportions of contributions from specific authors are considered. The metric is able to account for a significant portion of the observed heterogeneity between studies included in meta-analyses. A meta-regression between observed effect measures and author bias values revealed that higher levels of author bias were associated with higher effect measures while lower author bias was evident for studies with lower effect measures. Furthermore, the software’s capabilities to analyse authorship contributions and produce scientometric plots was able to reveal distinct patterns in both the temporal and geographic distributions of publications, which may relate to any evident publication bias. Thus, ABCal can aid researchers in gaining a deeper understanding of the research landscape and assist in identifying both key contributors and holistic research trends.

## Introduction

Scientific studies based on the empirical method contribute dozens of new publications pertaining to a central hypothesis under investigation on an annual basis. Over time, however, it may become apparent that the evidence in support of, or opposing, fundamental views within a discipline may not be unanimous (Le Clercq, Bazzi, et al. 2023) and can confound the overall interpretation of primary findings. As existing evidence serves as the foundation for informing the direction of prospective studies, however, scientists are often faced with the difficult task of reading and interpreting the literature to derive central tenets for their subject or discipline (Boell and Cecez-Kecmanovic 2010; Webster and Watson 2002). One convenient method to synthesise and assess the existing evidence is through systematic review and meta-analysis.

Systematic reviews were first developed as a tool for the synthesis of evidence for causality (le Clercq et al. 2016) or treatment (Honvo et al. 2019) in medical research and can be defined as: “a review using a systematic method to summarize evidence on questions with a detailed and comprehensive plan of study” (Tawfik et al. 2019). Thus, a systematic review seeks to identify and critically evaluate all those studies (Dickersin et al. 1994) pertaining to a specific research question for the purposes of deriving conclusions from the full body of evidence rather than relying on individual studies alone. Furthermore, systematic reviews attempt to standardise the methods (Moher et al. 2010; O’Dea et al. 2021) used to identify and screen studies in a way that is comprehensive, transparent, and above all reproducible and can serve as independent studies (Kraus et al. 2022). This avoids some of the pitfalls and biases that could influence narrative reviews (Pae 2015; Tawfik et al. 2019). Another advantage of systematic reviews is the possibility to perform a meta-analysis and scientometric assessment of the included studies (Nakagawa et al. 2023). This is done using primary reported statistics to derive the effect size (Cohen 1988) or treatment effect (TE), and variance or standard error (SETE), of the measured outcome. This facilitates between-study comparisons and enables the calculation of a pooled effect through a fixed- or random effects model (Borenstein et al. 2010). The pooled effect, therefor, serves as a quantitative measure of the total evidence.

Meta-analysis is not without possible confounders: several factors could contribute to between study differences, called heterogeneity (Higgins and Thompson 2002), or be a source of bias (Felson 1992; Sterne et al. 2001). Heterogeneity is expressed by two statistics, the heterogeneity measure (*I*^2^) and the between-study variance or tau-squared (*τ* ^2^). The *I*^2^ measure expresses the percentage of total variance in the effect sizes that is explained by between-study variance. The *τ* ^2^ approximates between-study variances but is reliant upon the specific effect sizes and needs to be quantitated by a P-value (Higgins 2008). As heterogeneity could potentially reduce the ability to compare or combine the outcomes from all studies that meet inclusion criteria, it is critical for authors to identify possible sources of heterogeneity and attempt to account for them.

Common factors that contribute to heterogeneity include sample size, quantitative method, and study population. Two approaches can be used to account for these variables: meta-analysis with subgroups and meta-regression of factors. The first, meta-analysis with subgroups, determines if effect sizes and their corresponding variances differ between subgroups (Borenstein and Higgins 2013). In the case of study populations representing different species, as may be the case in reviews in animal sciences or ecology, several methods have also been developed to account for phylogeny (Chamberlain et al. 2012) and taxonomy by performing a phylogenetic meta-analysis (Adams 2008; Lajeunesse 2009). The second approach, meta regression, determines if a significant part of the heterogeneity can be accounted for my individual study attributes, which may be more useful for continuous variables such as sample size (Baker et al. 2009).

The other, and perhaps more difficult, task is to identify and quantify potential sources of bias (Boutron et al. 2019; Felson 1992; Sterne et al. 2001). The most (Bouyssou and Marchant 2016; Perianes-Rodriguez et al. 2016; Zhou and Leydesdorff 2010) established form is publication bias (Lortie et al. 2007; Møller and Jennions 2001; Thornton and Lee 2000). This form of bias is detected through funnel-plots or through weighed linear models and focusses on detecting small study effects (Egger et al. 1997; Sterne et al. 2001). This includes the absence of studies that have smaller sample sizes, possibly due to the difficulties associated with the peer-review and publishing of such studies, as well as the inclusion of studies with small sample sizes and very low variance. There is, however, another related source of bias that has received little to no recognition thus far–bias from the overrepresentation of studies from specific authors (Ausloos 2013); hereafter, author bias.

Author bias, when not accounted for, has the inherent ability to skew the overview and interpretation of the literature in several ways. In the first, a specific view may be held by a particular group of authors who publish at a much higher frequency than other scientists in their field (Lortie et al. 2007), resulting in many publications in support of a view from a narrow pool of authors. This may create the illusion that opinions reported in their papers represent a majority consensus even when few independent studies support their claims. Secondly, views derived from primary findings based on a novel and ‘in-house’ method may not be fully reproducible if no independent studies exist where other authors repeated and confirmed the validity of such methods. This can further be confounded by the fact that negative or disconfirming results are often published at a delay (Boutron et al. 2019) or in less prominent journals (Leimu and Koricheva 2004). Lastly, a majority of studies may have been conducted in a specific country, region (Collyer 2018), and context (Fohringer et al. 2022) which–in cases where study populations may vary significantly between regions–may result in interpretations and generalisations that aren’t universal. This makes the scientometric analysis of studies by author, year, and location, critical in providing an appraisal of the literature.

At present, the most common methods used to access author contributions are the use of fractional citation counts (Bouyssou and Marchant 2016; Perianes-Rodriguez et al. 2016; Zhou and Leydesdorff 2010). This method emerged as a practical approach in response to the various complexities of assessing the contribution levels within scholarly works attributed to specific authors. While this method has proven useful in illuminating prominent contributors in differing fields (Bedru et al. 2023; Small and Garfield 1985) and in citation network analyses (Perianes-Rodriguez et al. 2016), no clear link has been made between scores for individual authors from fractional counting and bias introduced in reviews from contribution levels. Furthermore, many of the methods that have been described still lack available software that implements the method (Bedru et al. 2023), or are available with very limited functionality (Keirstead 2016; Kozlowski 2019). To address the current need to quantitate author contribution levels as a measure of bias, and perform scientometric checks on publication year and location, ABCal (version 1.0.2) was created to compute author bias and plot scientometric aspects of studies included in systematic reviews and meta-analyses. In this paper, a full description is given of how the author bias metric is computed along with examples of how ABCal can be used to evaluate potential sources of bias using real data from two datasets from a recent systematic review (Le Clercq et al. 2023; Le Clercq, Kotzé, et al. 2023).

## Methods

### Author Bias metric

To assess relevant attributes of included studies, ABCal includes a new metric referred to as “author bias” to provide a quantitative measure for the possible effect of over-represented authors introducing bias to the overall interpretation of the literature. This measure is derived in several steps. The theoretical framework for the metric is described in Fig S1. First, the full list of authors for all included studies (*L*_*All*_) is used to determine the total number of times the name of a specific author (*n*_*Author*_) occurs when iterating through each position in the list from *i* = 1 to #*L*_*All*_ (Equation 1).

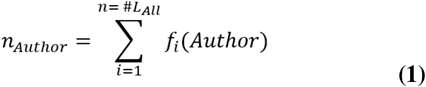

This value is then divided by the total number of authors in the list (#*L*_*All*_) which provides the individual author bias (*AB*_*Ind*_), as the proportion of total authorship contributions that belong to individuals (Equation 2).

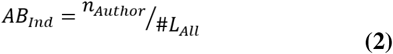

Next the bias derived from authorship is calculated per study (*AB*_*Study*_) by adding the individual bias values for each author in the author list for a specific study (*L*_*Paper*_), per Equation 3.

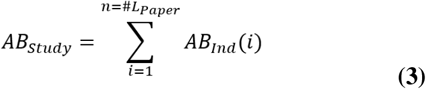

To facilitate the interpretation of these values, the final steps are calibration (Equation 4), by dividing total bias per study (*AB*_*Study*_), by the number of authors per paper (#*L*_*Paper*_).

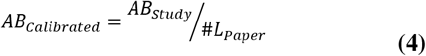

These are further categorised by assessing the distribution of author bias values by identifying those studies that fall in the bottom, middle, and upper third range, or percentiles (Equation 5) of thirty-three, to assign bias status as being low, medium, or high based on the calculated quantiles.

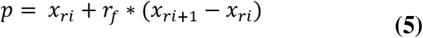

ABCal also provides some functionality to assess the normality of the author bias values using three approaches: a Shapiro-Wilk test (Shapiro and Wilk 1965), a Quantile–Quantile (QQ) plot (Wilk and Gnanadesikan 1968), and a histogram of distributions. The plotting sub-menu also provides an option for plotting the distribution of z-score transformed bias values which is useful in comparing distributions for different meta-analysis datasets.

The performance of the author bias metric was assessed for both validity and reliability (Cohen et al. 2017; Hammersley 1987). Validity, in this context, refers to the ability of the metric to accurately measure the intended attribute. This was verified by comparing those studies for which a higher author bias value was computed to whether the authors listed on the paper ranked within the top 10 contributing authors for the field. Agreement was measured in *R 4*.*0*.*6* (R Core Team 2020) using Cohen’s kappa (Cohen 1960) with the *vcd 1*.*4-11* package (Meyer et al. 2023). Reliability or repeatability of the measure was assessed by comparing the paper level calibrated author bias levels coded as low (1), medium (2), or high (3), for two different datasets (Le Clercq, Grobler, et al. 2023). This was done by assessing their independent validity as well as internal consistency between the distributions using Cronbach’s alpha (Cronbach 1951) with the *ltm 1*.*2-0* package (Rizopoulos 2006).

### Implementation

ABCal, *version 1*.*0*.*2* (Le Clercq 2023), was scripted in the Spyder 5 IDE using the PYTHON 3 language (Python Team 2021) and should be compatible with all versions upward of *version 3*.*6*. The list of packages that form part of the dependencies is provided on the GitHub repository and within the README file, along with detailed instructions for the download and installation. Dependencies include the use of several core PYTHON based libraries such as *NumPy 1*.*20*.*1* and *pandas 1*.*2*.*4* to handle input files and mould data structures for analyses (Harris et al. 2020; McKinney 2010). Other dependencies include packages for statistical analyses, such as *SciPy 1*.*6*.*2* (Virtanen et al. 2020) and *statsmodels 0*.*12*.*2* (Seabold and Perktold 2010), and packages for graphical plotting, such as *matplotlib 3*.*3*.*4* (Barrett et al. 2005) and *GeoPy 2*.*3*.*0* (Lopez Gonzalez-Nieto et al. 2020). ABCal further uses the plotting functionality implemented in *folium 0*.*14*.*0* with selected functions from the IO tools, for input and output, as well as the PYTHON Image Library (*PIL 10*.*0*.*0*). Menu options (detailed in ‘Usage’ section) provide the utilities to calculate author bias, test the distributions of author bias values, and generate several scientometric plots. Scientometric plotting options include the ability to plot publications by the top contributing authors (to identify and visualise the extent to which top authors may skew overall interpretation), by year, and by location.

### Input and Output file formats

All input files used by ABCal are in the standard comma separated value (CSV) format. For most functions the first column of these files should contain the heading “Paper” with the studies listed by name in the column e.g., “Le Clercq *et al*. (1987)”. To calculate the author bias, the CSV file should contain columns for each author, first to last, labelled with appropriate headings such as “Author1” etc. These columns should contain the last name and initials of each author associated with a specific paper e.g., “Le Clercq, L.S”. The function to compute author bias moves through several steps to perform each intermediate step to derive the values and provides intermediate output files with relevant measures for later steps or scientometric plotting, detailed under the usage section. These files provide the option to specify unique output file names and are stored as CSV files within the current working directory.

For most of the plotting options, either the CSV files generated from author bias computation or CSV files containing additional study attributes for plotting are used. An example of such a file to plot the distribution of publications by year, includes a file containing two columns with the headers “Paper” and “Year”, which should contain the study name as well as the year of publication. Another example is for the plotting of studies by location, where a CSV is required containing two columns with the headers “Paper” and “Location”. For this file, the full name of the country in which the study was conducted is required; if more than one location was included these should be listed on separate lines with the study name and second or third location. The output generated from plotting is saved with a standard name for the type of plot in the portable network graphic (PNG) format, with the exception of the location plot which is also stored in the interactive HTML format.

### Usage

To illustrate the usage of ABCal, two datasets (Le Clercq et al. 2023) generated as part of a recent systematic review on biomarkers for age in animals (Le Clercq, Kotzé, et al. 2023), comprising age models from included studies on the use of methylation (N = 41 studies, 60 models) and telomeres (N = 67 studies, 99 models) respectively, will be used. For each dataset, three input files were generated: one with the paper name and list of authors, a second with the paper name and publication date, and a third with the paper name and study location.

The first file was used to compute the author bias (example A). Once ABCal is initiated, the first option (a) is to calculate the total author bias per paper. Selecting this option initiated the function to perform the needed steps to do the calculation. A prompt appeared to specify the name of the file containing the author lists e.g., ‘Auth Meth.csv’. The first step generated a list of all authors along with the total counts of times the specific author appeared in an author list. This data was exported as the first file for output from the function and was saved as a CSV containing the number of publications per author. Next, the individual author bias was computed by determining the fraction of total authorship contributions per author. These values were stored in the second output and contained the author names and their associated individual bias. The final step computed the total author bias per paper by adding the individual bias value of each author associated with the author list for a paper. This data was saved as the third output and contained the data used to compute total bias per paper. As an additional step, and to assist in the interpretation of values, the second menu option (b), which takes the final output file from the first option, was used to calibrate the bias value by dividing the total bias by the number of authors per paper. The newly calibrated values were exported as the fourth output file. Furthermore, the third menu option (c) was used for testing the distributions for normality and the fourth menu option (d) was used to get the upper, middle, and lower third quantiles of the author bias distributions.

Hereafter, the author bias values and their respective levels were incorporated into two meta-analyses as part of a review (Le Clercq, Kotzé, et al. 2023). The meta-analysis was done in RStudio 1.4.1106 (RStudio Team 2021), running R 4.0.5 (R Core Team 2020) with the package meta 5.5-0 (Harrer et al. 2021; Schwarzer et al. 2015). A meta-regression was done between the random effects model and author bias as a predictor of heterogeneity for a functional test of validity. The results were visualised using a bubble plot implemented in the metafor 3.8-0 package (Viechtbauer 2010) with grouping based on the three quantiles. Furthermore, potential publication bias as measured by funnel plot asymmetry was also assessed using the Egger’s test (Egger et al. 1997) as implemented in metafor 3.8-0, also plotting the relationship between the standardized measured effect and the inverse of the standard error.

Scientometric plotting capabilities of ABCal, accessed via a submenu when selecting option e, were illustrated in example B for methylation studies and example C for telomere studies. For the first option (a) from the submenu, the second file with two columns for ‘Paper’ and ‘Year’ was used to plot the total number of publications per year. The first file given as output from the author bias computation steps, containing the authorship counts, was used for the second option (b) to plot the number of contributions from the top ten contributing authors. Lastly, the third file containing two columns for ‘Paper’ and ‘Location’ was used to plot a choropleth map of the geographical distribution for study locations using the third option (c) from the submenu.

## Results

### Example A – Author Bias Computation

Author bias values per paper for methylation studies ranged from 0.0024 to 0.0302 with a mean of 0.0124 (Table 1). The histogram plot of distributions (Fig. 1A) showed many values (N = 19) were well below the mean, skewing the distribution left, with a moderate number of studies (N = 10) falling in and around the mean and few studies (N = 12) having higher values. The position for setting the first (Q1) and third (Q3) quartiles were 0.004 and 0.018 respectively, with 14 studies classified as low, 13 as medium, and 14 as high. The box plot of z-score transformed values (Fig. 1B), to express the bias values in terms of standard deviations from the median, showed that the median was low (approximately -0.094) with most studies (95 percent) being evenly distributed around the median. A few studies had higher values; however, they still fell within two standard deviations of the median and no clear outliers were detected. The overall distributions were found to not be normally distributed (Table 1; P < 0.01). Tests for validity by means of Cohen’s kappa showed high levels of agreement (Table 2; *κ* = 0.94, P < 0.01) between studies ranked as having medium to high risk of bias as compared to the list of top contributing authors.

**Table 1.** Summary of characteristics of calibrated author bias values. Values are reported for both the calibrated author bias (AB) values as well as the Z-Score transformed values. The mean, standard error of the mean (SE), minimum (Min), maximum (Max) and positions of the cut-off points for the lower (Q1) and upper (Q3) third percentiles are given. The results for the normality test, tested using the Shapiro-Wilk test, are also given along with the significance. The calibrated author bias for both datasets was not normally distributed (P < 0.01). The number of studies identified by level as having either low, medium, or high risk of bias are also indicated.

|  | Mean | SE | Min | Max | Q1 | Q3 | Shapiro-Wilk |
| --- | --- | --- | --- | --- | --- | --- | --- |
| **Methylation:** |  |  |  |  |  |  |  |
| *Author Bias* | 0.014 | 0.001 | 0.002 | 0.03 | 0.004 | 0.018 | 0.869 ($P < 0.01$) |
| *Z-Score* | $-1.08 \times 10^{16}$ | 0.16 | -1.07 | 1.9 | -1 | 0.75 | |
| (1) *Low* | 14 |  |  |  |  |  |  |
| (2) *Medium* | 13 |  |  |  |  |  |  |
| (3) *High* | 14 |  |  |  |  |  |  |
| **Telomeres:** |  |  |  |  |  |  |  |
| *Author Bias* | 0.004 | 0.0001 | 0.002 | 0.012 | 0.003 | 0.004 | 0.780 ($P < 0.01$) |
| *Z-Score* | $-2.10 \times 10^{16}$ | 0.12 | -0.79 | 4.13 | -0.79 | 0.44 | |
| (1) *Low* | 23 |  |  |  |  |  |  |
| (2) *Medium* | 21 |  |  |  |  |  |  |
| (3) *High* | 23 |  |  |  |  |  |  |

**Table 2.**
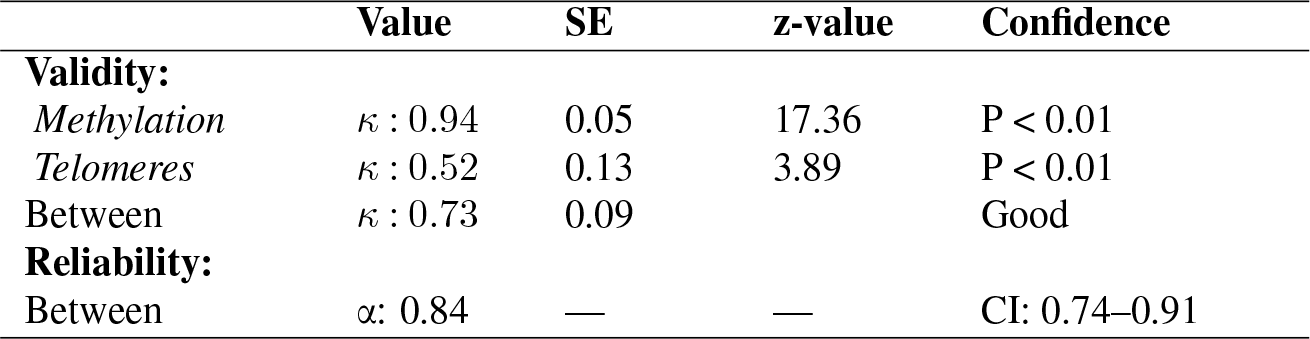
Results from tests for validity, by mean of Cohen’s kappa (κ), as well as reliability, by means of Cronbach’s alpha (α), for the Author Bias metric. Validity was calculated for both the methylation and telomere dataset by assessing the agreement between bias rank and authors listed as the top ten contributors. The overall validity was determined by taking the average for individual results. Agreement for ranking between datasets was used to assess between study reliability of the metric. The confidence of the calculated values was assessed by either probability (P < 0.01) of a z-test or the 98% confidence interval (CI).

**Fig. 1.**
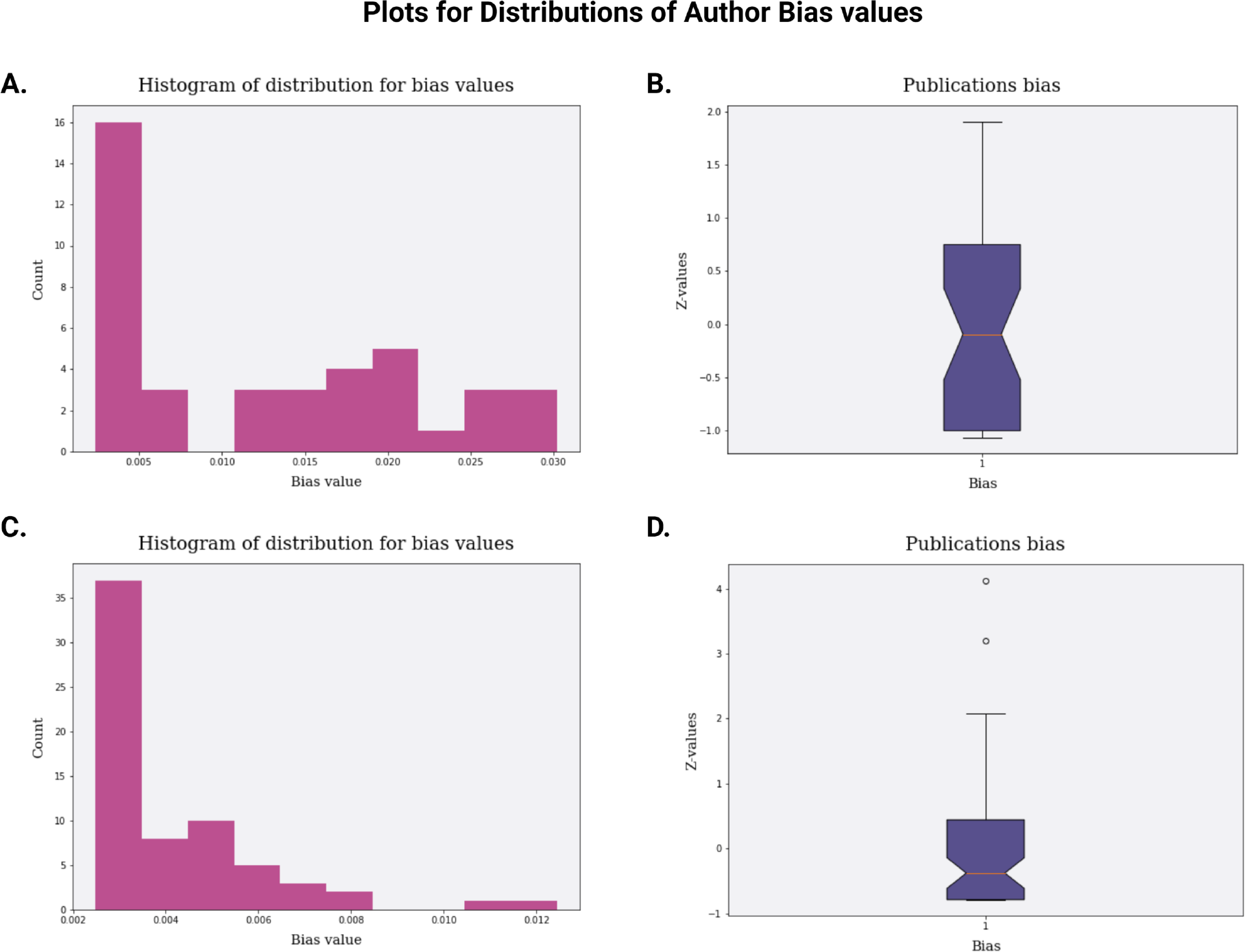
Plots for the distributions of author bias values. A. Histogram for the calibrated author bias values for papers in the methylation dataset indicating many studies (N = 16) with low values, skewing the distribution left, with a moderate number of studies (N = 4-6) around the median and a similar number of studies (N = 3-6) with high values. B. Box plot for the Z-score transformed author bias values for papers in the methylation dataset indicating most studies are evenly distributed around the median (orange line) with few studies more than one standard deviation from the median, and a small number of studies in the upper range without appearing as outliers. C. Histogram for the calibrated author bias values for papers in the telomere dataset indicating many studies (N = 25) with low values, skewing the distribution left, with a moderate number of studies (N = 5-10) around the median and two studies with higher values. D. Box plot for the Z-score transformed author bias values for papers in the telomere dataset indicating most studies were slightly above the median (orange line) with most studies within one standard deviation from the median, and a small number of studies in the upper range and two outliers. (image created in BioRender.com)

Author bias values for telomere studies ranged from 0.0024 to 0.0124 with a mean of 0.0040 (Table 1). The position for setting the first (Q1) and third (Q3) quartiles were 0.003 and 0.004 respectively, with 23 studies classified as low, 21 as medium, and 23 as high. The values followed a similar pattern to that observed for methylation studies (Fig. 1C), with a large number of studies (N = 48) falling below the mean. Most of the remaining studies followed a near bell shape around the mean, with several (N = 9) having higher values. The box plot (Fig. 1D) showed a slightly higher number of studies fell above the median, while most studies were still within one standard deviation of the median. Several studies had values between one and two standard deviations of the median, with two studies that were more than two standard deviations from the median and thus detected as outliers. Once more, the distributions were found to not be normally distributed (Table 1; P < 0.01). Tests for validity showed a moderate, yet significant, level of agreement (Table 2; *κ* = 0.52, P < 0.01) and an average validity between datasets of 0.73. The reliability tests between datasets also showed a significant (Table 2; α = 0.84, CI: 0.74–0.91) level of reproducibility for levelled classification of studies.

A meta-regression between observed effect measures and author bias values revealed that author bias values were able to account for a significant portion of the observed heterogeneity between studies included in meta-analyses. These relationships are illustrated as bubble plots in Fig. 2. For methylation studies, a strong relationship was observed (P < 0.01) with author bias values accounting for approximately 23 percent (*R*^2^ = 0.23) of the study heterogeneity. For telomere studies, a slightly weaker relationship was observed (P < 0.02) with author bias values accounting for 6 percent (*R*^2^ = 0.06) of the study heterogeneity. In both instances, higher levels of author bias were associated with higher effect measures while lower author bias was evident for studies with lower effect measures. Tests for publication bias (Fig. S2) detected significant funnel plot asymmetry (P < 0.05) indicative of possible publication bias. Statistical methods to address publication bias, such as “trim- and-fill” or linear modelling of a fixed-effect model with factorisation, did not significantly alter the overall interpretations from the comparisons (data not shown).

**Fig. 2.**
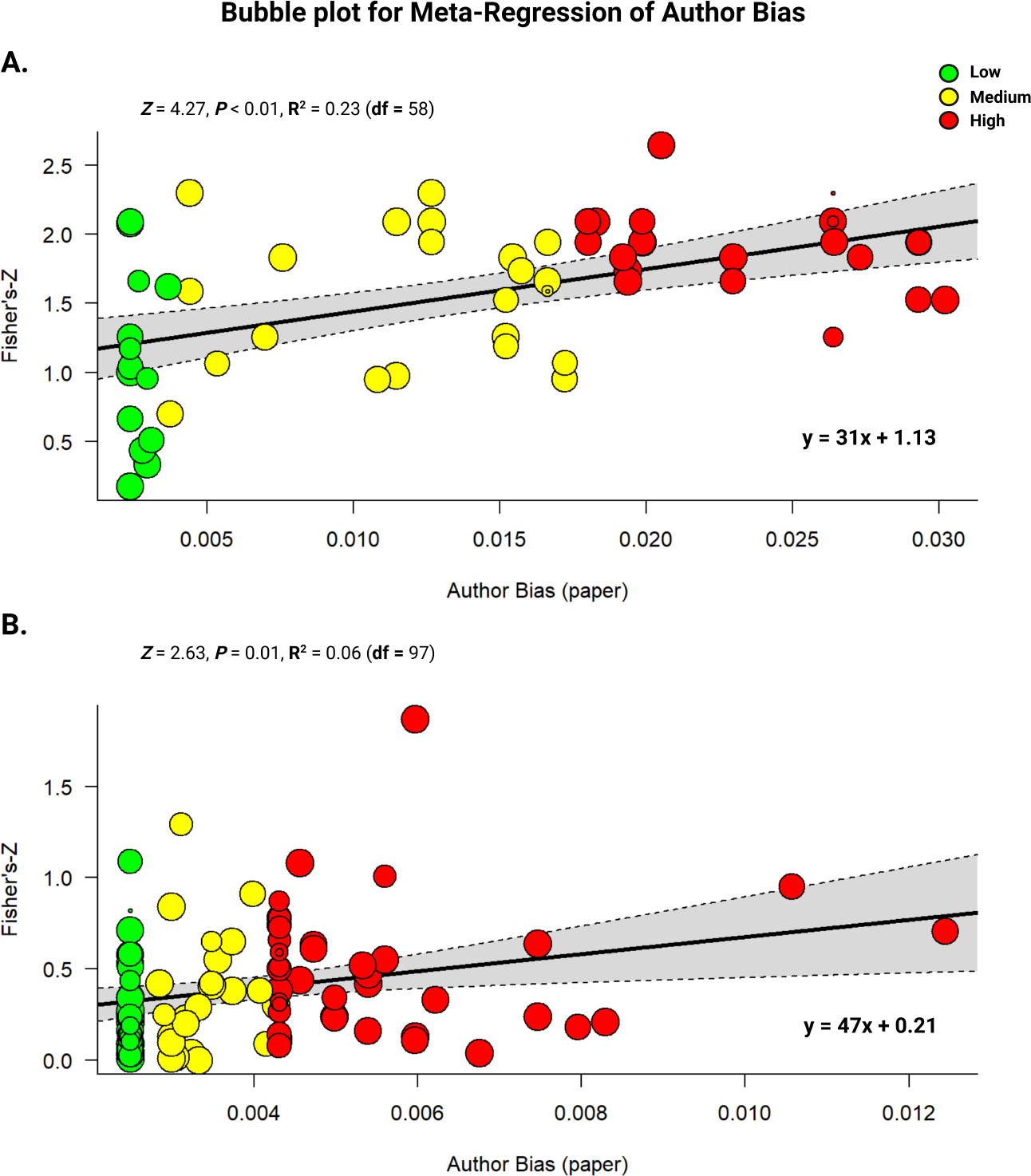
Bubble plots for the meta-regression of author bias values as a predictor of heterogeneity in the meta-analyses. The Fisher’s-Z values (y-axis) were plotted against the calibrated author bias values per paper (x-axis) and studies colour coded according to the quantiles within which they fell and were classified: low (green), medium (yellow), or high (red). The linear equations for the regressions are indicated in the bottom right of each plot. A. Meta-regression performed on the methylation dataset found a significant association between effect measures and author bias values (P < 0.01), accounting for 23 percent of the observed heterogeneity. B. Meta-regression for the telomere dataset found a moderate association (P < 0.02), accounting for only 6 percent of the heterogeneity. (image created in BioRender.com)

### Example B – A Meta-Analysis of Methylation Studies

Scientometric assessment of studies included in the methylation dataset was done by plotting three attributes: top contributing authors, publications by year, and publications by location. For publications per author, the number of publications contributed by the top contributing authors (specified as ten) ranged from 3 contributions to a total of 24 contributions (Fig. 3A). Five authors, including Zhang, contributed 3 papers each, respectively. The top three contributing authors, identified as Horvath, Haghani, and Zoller, contributed to approximately half (21-24 out of 41) of the included studies. The bar plot for publication by year (Fig. 4A) revealed the first studies were published in 2014 (Polanowski et al. 2014) with an annual increase leading to 15 publications in 2021 (Bors et al. 2021; Mayne et al. 2021; Robeck et al. 2021; Wilkinson et al. 2021) and several recent studies (Horvath, Haghani, Peng, et al. 2022; Horvath, Haghani, Zoller, et al. 2022; Robeck et al. 2023). The choropleth map (Fig. 4B), showing study locations, showed the number of studies per country ranged from one study (green) to more than twenty studies (red). Several countries, shown in white, were completely data deficient. The overall distribution showed that most studies emanated from the Northern hemisphere, principally from North America (N > 20) and Europe, with Australia (N > 5) representing the country with the most publications in the global South.

**Fig. 3.**
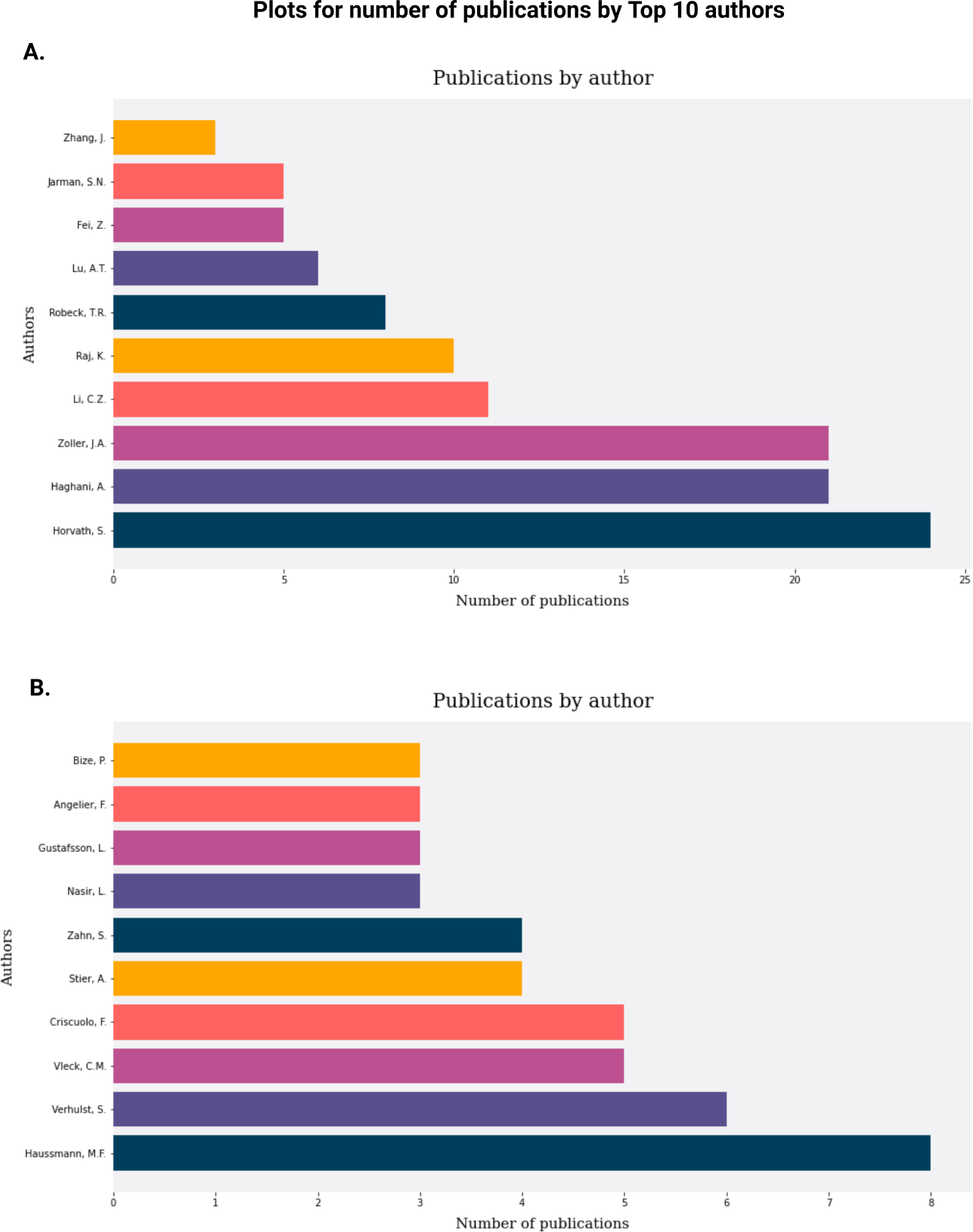
Scientometric plots for the number of publications attributed to the top 10 contributing authors. A. For methylation studies, the number of publications attributed to top contributing authors ranged from 4 contributions to as many as 24 contributions: with the top 3 contributing to a half (50%) of the included studies. B. For telomere studies, the number of publications attributed to the top contributing authors ranged from 3 to 8 contributions: here, the top 3 contributing authors made up approximately a tenth (10%) of the total studies. (image created in BioRender.com)

**Fig. 4.**
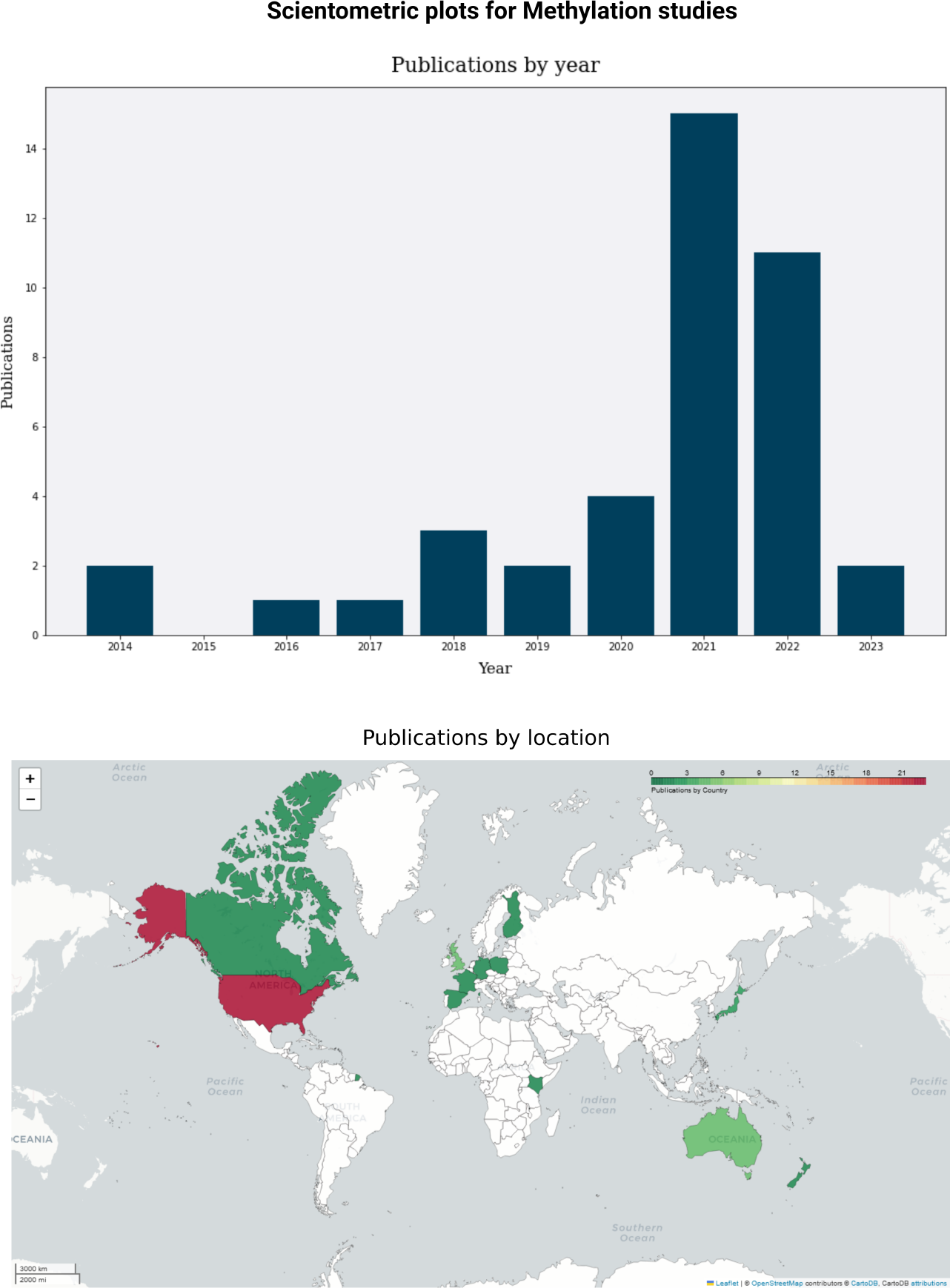
Scientometric plots for methylation studies, generated with ABCal. A. Bar plot for publication by year, indicating the first studies published in 2014 with an annual increase leading to 15 publications in 2021. B. Choropleth map showing study locations. The density gradient plots the number of studies per country ranging from one study (green) to more than twenty studies (red); countries in white are data deficient. The overall distribution shows most studies are from the Northern hemisphere, principally from North America and Europe, as well as Australia. (image created in BioRender.com)

### Example C – A Meta-Analysis of Telomere Studies

The same scientometric plots were also generated for the telomere dataset. For publications per author, contributions by the top ten authors ranged from 3 contributions to a total of 8 contributions (Fig. 3B). Several authors contributed 3 papers while the top three authors, identified as Haussmann, Verhulst, Vleck, and Criscuolo, each contributed between 5 and 8 studies. This only accounted for about 10 percent of the included studies (5-8 out of 68). Publications by year, given as a bar plot (Fig. 5A), indicated the first included studies were published circa 2002 (Brümmendorf et al. 2002; Haussmann and Vleck 2002) with frequent subsequent publications, around 2-3 studies annually, and an increase seen after 2012 (Fick et al. 2012; Plot et al. 2012) with several spikes in 2017 (Cerchiara et al. 2017; Kirby et al. 2017; Ujvari et al. 2017), 2020 (Bauch et al. 2020; Burraco et al. 2020; Cherdsukjai et al. 2020), and 2021 (Molbert et al. 2021; Vernasco et al. 2021). The maximum number for spikes ranged between 6 to 8 publications. Choropleth mapping of study locations as publications per country (Fig. 5B) ranged from one study (green) to ten studies (red); data deficient countries are indicated in white. The distribution showed most studies originated from the Northern hemisphere, principally from North America (N > 9). The most represented country from the Southern hemisphere was Australia (N > 6).

**Fig. 5.**
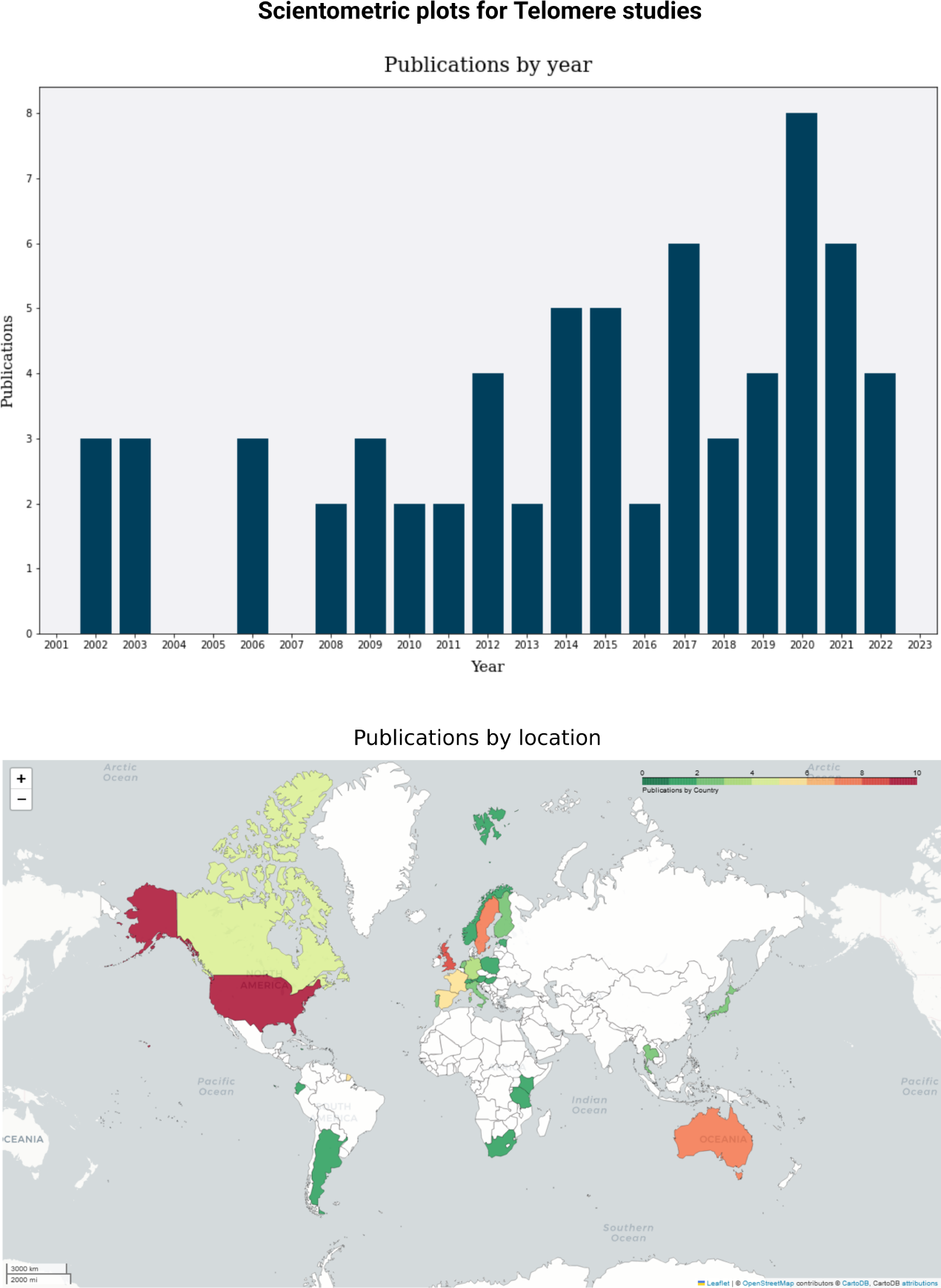
Scientometric plots for telomere studies, generated with ABCal. A. Bar plot for publications by year, showing the first included studies were published in 2002 with constant publication of 2-3 studies per year and increasing from 2012 with several spikes in 2017, 2020, and 2021 to between 6-8 publications. B. Choropleth map showing study locations. The density gradient plots the number of studies per country ranging from one study (green) to ten studies (red); countries in white are data deficient. The distribution shows most studies are from the Northern hemisphere, principally from North America, as well as Australia. (image created in BioRender.com)

## Discussion

This original paper presents the methods and results of a research study using a newly developed software tool called ABCal. The tool is implemented in Python and is designed to analyse scientometric data from various studies when conducting systematic reviews and meta-analyses. The primary focus of the study was to compute and assess author bias in the literature, providing a quantitative measure for the possible effect of overrepresented authors introducing bias to the overall interpretation of the literature. The computed author bias values provide a quantitative measure of author influence on the interpretation of the studies. Furthermore, scientometric plots provided valuable insights into the trends and distribution of publications over time and geographic locations.

The distribution of author bias values, as shown in histograms and box plots, showed marginal differences in the raw values but was conserved between the two datasets when using z-score transformed values. This highlighted similar spreads for the distribution with low to medium bias for most studies and a smaller number of studies exhibiting higher bias levels. As such, author bias values were able to identify, in a quantitative manner, the overrepresentation of some authors in both meta-analytic datasets. When combined with scientometric plots of author contributions, it became clear that both datasets contain a significant number of authors who have contributed a larger number of studies than others. This makes the proposed author bias metric useful in addressing the account for such bias when doing reviews (Felson 1992; Knobloch et al. 2011).

The author bias metric is organically related to fractional citation counting (Zhou and Leydesdorff 2010), an emerging ‘golden standard’ when comparing author contribution levels. This is due to similarities between calculations used in the initial steps that count the total number of times an author appears in the list of authors, which is divided by the total number of authors in the list (Bouyssou and Marchant 2016). Considering, however, that this is only done within the context of studies included in a meta-analysis instead of the full reference list—in this instance the new metric represents a special use case of fractional counting. Subsequent steps sum the fractional count for individual authors per included study and divides the total by the number of authors per paper to derive a paper-level value for author bias. Furthermore, rather than relying on raw values, ABCal provides the option to convert between raw values and z-scores as well as three levels of interpretation: low, medium, and high. This facilitates cross-discipline use of the calibrated author bias metric as several cultural differences may exist between fields in terms of publication and citation behaviour (Bornmann and Daniel 2008; Zhou and Leydesdorff 2010). At present ABCal, and the novel author bias metric, also provides enhanced utility in comparison to existing options (Bedru et al. 2023; Keirstead 2016; Kozlowski 2019). For example, ABCal uses provided information and does not rely on the indexing of papers on a specific database (Kozlowski 2019) or the existence of author profiles on a specific platform (Keirstead 2016). This is particularly important for included studies from publishers that don’t index their articles on all databases or when including preprints from e.g., bioRxiv. ABCal is also freely available to the community for implementation in other studies while several similar algorithms are not (Bedru et al. 2023).

Any new metric is, however, subject to benchmarking through tests of validity and reliability (Cohen et al. 2017; Hammersley 1987). Validity, as measured by agreement between medium to high bias studies and the list of top authors using Cohen’s kappa (Cohen 1960), showed moderate to high agreement levels that are generally well suited given the application (Altman 1990). Reliability, as measured by Cronbach’s alpha, found that the author bias metric was able to partition studies into low, medium, and high bias with a high degree of consistency between datasets. Furthermore, the functional validity was assessed by meta-regression for which the results indicated a significant association between author bias and observed effect measures in the meta-analyses. More specifically, higher author bias values were associated with higher effect measures, while lower bias was evident in studies with lower effect measures. This also makes author bias values utile in understanding how author prominence (Cassey et al. 2004), from higher contributions to the field, may interact with reported effect sizes to account for part of the heterogeneity that exists between studies as well as publication bias (Baker et al. 2009).

The presence of publication bias, as revealed by tests of funnel plot asymmetry (Møller and Jennions 2001), suggests the possibility of selective publication in the literature (Boutron et al. 2019). This is typically attributed to small study effects such as the exclusion of studies with smaller sample sizes, even when sample sizes are sufficient for adequate statistical power of a given test (Cohen 1988; Faul et al. 2009; Kang 2021). The concentration of studies in certain regions as seen by geographic mapping of study locations, however, indicates a potential research trend of fewer or missing studies from the global South, as previously suggested (Collyer 2018), and evidence that research in ecology or animal science may not follow a truly global distribution (Martin et al. 2012). Considering researchers from lower income countries may conduct research on a smaller scale for economic reasons, it is feasible that the existing evidence of publication bias is due to the lack of studies from the Southern hemisphere in the literature.

It’s important to acknowledge the limitations of the study. The analysis relies on the accuracy and completeness of the input data (Knobloch et al. 2011; O’Dea et al. 2021), and certain assumptions might have been made during the calculation of author bias. Additionally, the analysis is limited to the specific datasets related to biomarkers for age in animals, and generalization to other research fields might require further investigation. Future work can focus on expanding the application of ABCal to different research areas and datasets to validate its effectiveness and robustness across various domains. Additionally, efforts can be made to address potential limitations and explore enhancements to the tool’s functionalities to meet the evolving needs of scientometric analysis in the ecology research community, particularly when conducting systematic reviews and meta-analyses.

## Conclusion

Overall, ABCal proves to be a useful tool for scientometric analysis, offering valuable information to researchers in assessing the impact of authors and potential biases in the literature. The software’s capabilities to analyse authorship contributions and produce scientometric plots can aid researchers in gaining a deeper understanding of the research landscape and identifying key contributors and research trends.

## Acknowledgements

All images were edited for publication in BioRender.com with publication licenses granted under the Academic plan. The author would like to thank Dr Desiré Lee Dalton, and Professors Paul Grobler and Antoinette Kotzé, for their support and advice. Open access provided by the University of the Free State. This work is based on the research supported wholly/in part by the National Research Foundation (Grant Number: 112062), South Africa.

## Data availability

The custom Python script for *ABCal version 1*.*0*.*2* is available for download for installation from source code on GitHub (https://github.com/LSLeClercq/ABCal), and includes example files used for testing. Data used in this paper were deposited online (https://doi.org/10.5281/zenodo.7091053) and recently published (Le Clercq et al. 2023)..

## Supplementary Information

**Fig. S1.**
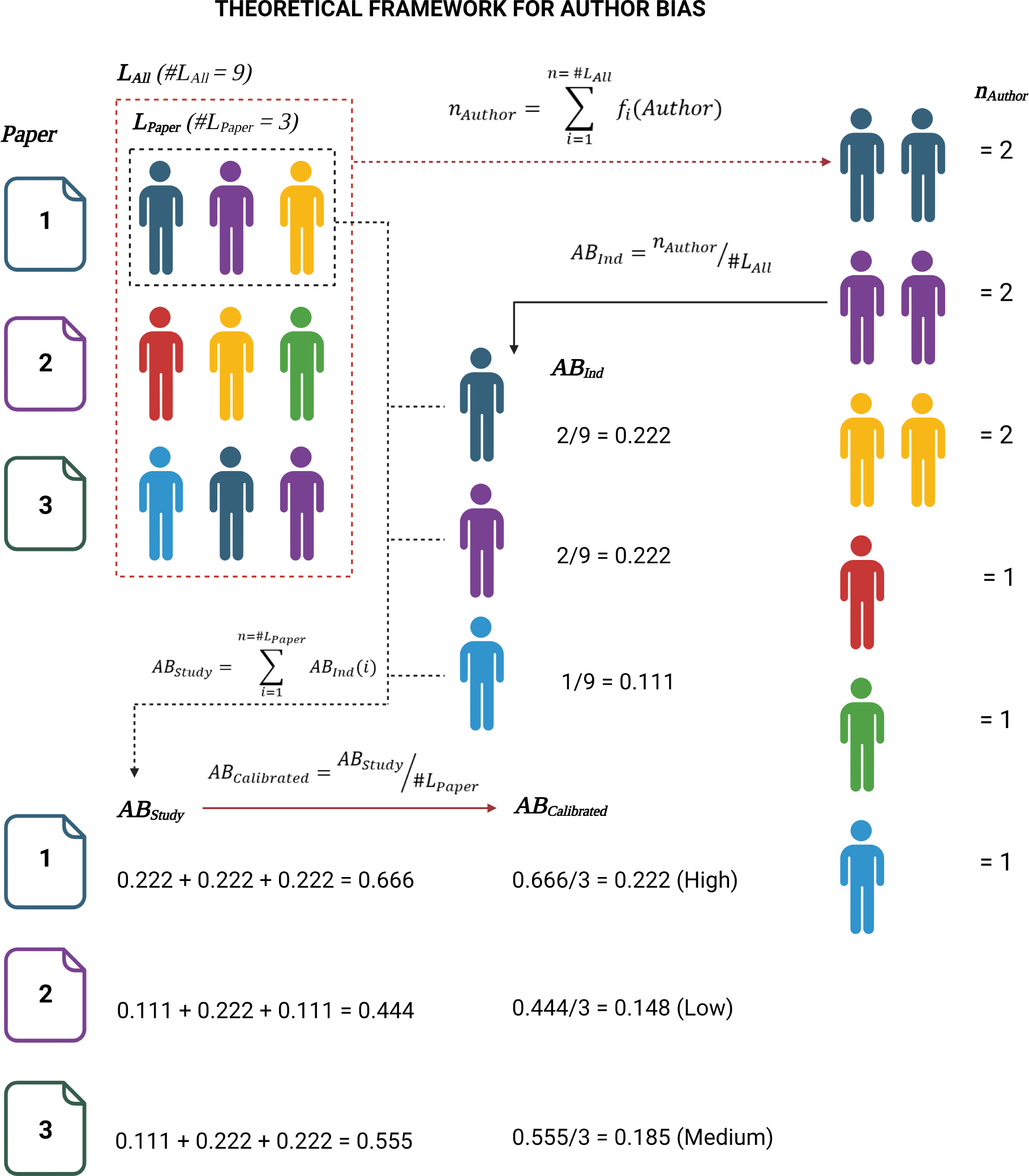
Theoretical framework for the calculation of author bias per paper. (image created in BioRender.com)

**Fig. S2.**
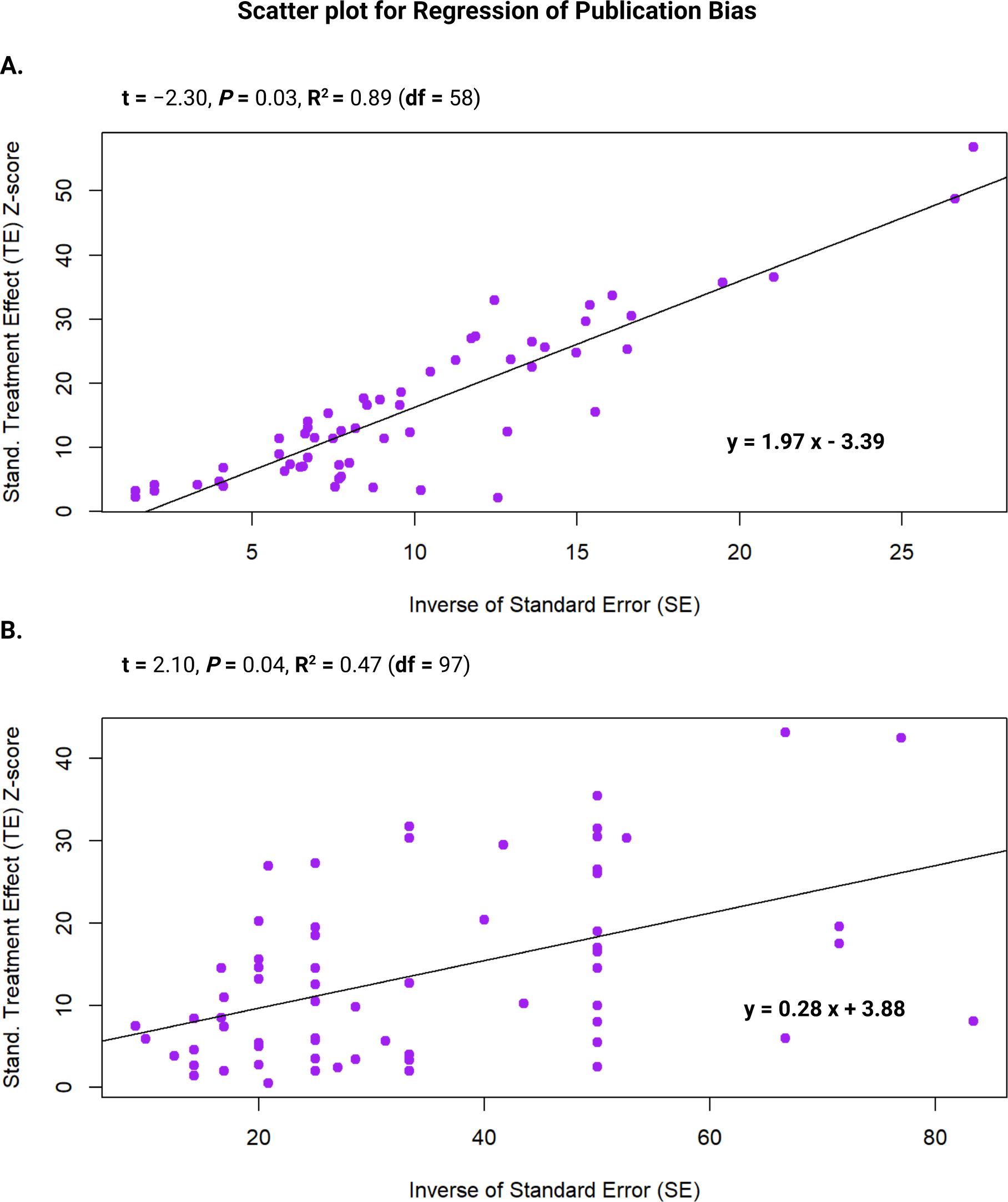
Scatter plots for the regression done to detect publication bias through funnel-plot asymmetry. The treatment effect (TE), Fisher’s-Z, as standardised to Z-scores (y-axis) were plotted against the inverse of the standard error (x-axis) for individual studies (purple). The fitted regression line from the Egger’s test is shown, along with the linear equation in the bottom right. A. For methylation studies, a strong correlation (*R*^2^ = 0.89) was detected along with significant indicators of potential publication bias (P < 0.05). B. For telomere studies, a slightly weaker correlation was detected (*R*^2^ = 0.43), however, there were still significant indicators (P < 0.05) for potential publication bias. (image created in BioRender.com)

